# The NHGRI-EBI GWAS Catalog: standards for reusability, sustainability and diversity

**DOI:** 10.1101/2024.10.23.619767

**Authors:** Maria Cerezo, Elliot Sollis, Yue Ji, Elizabeth Lewis, Ala Abid, Karatuğ Ozan Bircan, Peggy Hall, James Hayhurst, Sajo John, Abayomi Mosaku, Santhi Ramachandran, Amy Foreman, Arwa Ibrahim, James McLaughlin, Zoë Pendlington, Ray Stefancsik, Samuel A. Lambert, Aoife McMahon, Joannella Morales, Thomas Keane, Michael Inouye, Helen Parkinson, Laura W. Harris

## Abstract

The NHGRI-EBI GWAS Catalog serves as a vital resource for the genetic research community, providing access to the most comprehensive database of human GWAS results. Currently, it contains close to 7,000 publications for more than 15,000 traits, from which more than 625,000 lead associations have been curated. Additionally, 85,000 full genome-wide summary statistics datasets - containing association data for all variants in the analysis - are available for downstream analyses such as meta-analysis, fine-mapping, Mendelian randomisation or development of polygenic risk scores. As a centralised repository for GWAS results, the GWAS Catalog sets and implements standards for data submission and harmonisation, and encourages the use of consistent descriptors for traits, samples and methodologies. We share processes and vocabulary with the PGS Catalog, improving interoperability for a growing user group. Here, we describe the latest changes in data content, improvements in our user interface, and the implementation of the GWAS-SSF standard format for summary statistics. We address the challenges of handling the rapid increase in large-scale molecular quantitative trait GWAS and the need for sensitivity in the use of population and cohort descriptors while maintaining data interoperability and reusability.

## Introduction

Genome-Wide Association Studies (GWAS) have expanded our understanding of the genetic architecture of complex traits, moving beyond single-gene assumptions to the polygenic nature of most human traits and diseases. This has been particularly transformative for conditions like diabetes (1), cardiovascular disease (2) and psychiatric disorders (3) where multiple genetic variants, each with a small effect size, contribute to disease susceptibility. Unlike other methodologies, like candidate gene studies or fine mapping, which focus on preselected genes or regions based on prior knowledge, GWAS adopts a relatively unbiased approach, analysing assayed or imputed genetic markers distributed across the genome with fewer assumptions about which regions are likely to be involved.

The NHGRI-EBI GWAS Catalog (www.ebi.ac.uk/gwas) includes all human GWA studies, defined as analyses of >100,000 variants genome-wide, based on either array or sequence-based genotyping and including targeted arrays. It is the largest and most complete publicly available resource for Findable, Accessible, Interoperable and Reusable (FAIR) (4) GWAS data, containing 625,113 curated SNP-trait associations (p<1×10-5) linked to >15.5K traits as of 1st July 2024. These genetic associations derive from 6,921 publications and correspond to 108,850 separate GWAS analyses. Full genome-wide summary statistics datasets - containing association data for all variants analysed regardless of p-value - are available for download for 66% of all journal-published GWAS in the Catalog. Additional prepublished datasets (e.g. associated with a preprint, under submission to a journal or never intended for journal publication), contribute to the total of >85K datasets. Full summary statistics are either deposited directly by authors via our submission portal (www.ebi.ac.uk/gwas/deposition) or obtained by our curators from external resources where they have been made available under a permissive licence (CC0, https://creativecommons.org/public-domain/cc0/ or equivalent) and meet stringent validation criteria (https://www.ebi.ac.uk/gwas/docs/methods/summary-statistics). Submissions can be made prior to journal publication, and embargoed until publication or made available immediately. As of 1st July 2024, there were ∼13,000 prepublished summary statistics datasets publicly available, plus additional studies from 52 submissions under embargo. Direct submission from authors has resulted in a massive increase in data volume available in the Catalog (currently ∼56TB) and improved utility of data for users - for example, for Mendelian randomisation, the generation of polygenic scores (PGS), and fine mapping to identify causal variants. Data submission has also contributed to those publications gaining 80% more citations than those not sharing summary statistics (5).

Identification of publications with GWAS is done using LitSuggest (6), a tool which uses machine learning to classify the several thousand publications indexed in PubMed each week. Every run results in an average of 40 GWAS candidate papers which are then triaged for inclusion in the Catalog, of which 15 meet stringent eligibility criteria (https://www.ebi.ac.uk/gwas/docs/methods/criteria). Curation is prioritised for studies with summary statistics, with SNP-Trait associations, large well-powered studies, and those with non-European ancestry. This ensures that the most valuable data is made available to users quickly.

Interpretation and cross-study integration of GWAS results critically depend on the correct characterisation of the trait of interest and the individuals included in the analysis. All GWAS Catalog studies are annotated with standardised metadata during curation to provide context about the study design and enhance reusability. Trait information is mapped to terms from the Experimental Factor Ontology (EFO) (7), ensuring that traits are consistently annotated across studies and interoperable with other data sources. Population descriptors are mapped to the labels described in our ancestry framework (8).

The GWAS Catalog is recognised as a Global Biodata Core Resource (https://globalbiodata.org/what-we-do/global-core-biodata-resources/list-of-current-global-core-biodata-resources/) and Elixir Core Data Resource (https://elixir-europe.org/platforms/data/core-data-resources), acknowledging its importance to the international life science and biomedical research communities. Between July 2023–June 2024, the GWAS Catalog was accessed from 205 countries and had more than 300,000 unique website users and ∼3.4 million page views, a ∼50% and ∼40% increase respectively over the same period in 2021–2022, described in our last database update. It also functions as a data feed to other key resources, such as Open Targets Genetics (9), the AMP Knowledge Portals (https://kp4cd.org/) (10), MRC-IEU OpenGWAS (11), GeneCards (12), Ensembl (13) and others, serving different user communities and enabling integration of GWAS data with other data types. The GWAS Catalog works closely with its sibling resource the PGS Catalog (14), a FAIR database of polygenic scores, sharing processes and ensuring data interoperability - for example, in trait and sample description. Reciprocal links between the two resources allow users to identify polygenic scores that have been generated using data in the GWAS Catalog, and in the PGS Catalog to identify the GWAS source data that was used to develop a given polygenic score; ∼20% of PGS Catalog publications are linked to source data in the GWAS Catalog providing traceability for scores.

### Improved Data content

The lack of standardisation in GWAS summary statistics is a long-standing issue affecting the ability of data consumers to combine or interoperate datasets and apply software tools without significant manipulation of files. Following a series of community working groups (15), wider community feedback and iteration, we proposed a standard for summary statistics data and metadata (16). The GWAS summary statistics format (GWAS-SSF) standard was finalised and implemented in the GWAS Catalog in April 2023, with newly released software pipelines for validation, formatting and harmonisation to conform to the public schema. GWAS-SSF defines mandatory fields (chromosome, base pair location, effect allele, other allele, beta/OR/HR, standard error, effect allele frequency and p-value) and recommended fields (such as rsID, reference allele, confidence intervals) in the summary statistics file, as well as a standardised set of accompanying metadata enabling efficient and accurate data re-use.

The submission process has been designed to minimise barriers to submission and maximise accessibility via a simple and easy to generate format whilst maintaining a high quality of data by mandating key content that supports data re-use. A 33% increase in the number of submissions per month after the publication of the standard indicates that this format is both accepted by the community and feasible for submitters. All submissions flagged as GWAS-SSF have passed mandatory data and metadata validation, and of those, ∼74% of data and ∼85% of metadata include one or more of the additional recommended fields. This suggests we have been successful in our goal to improve the quality of summary statistics without placing onerous requirements on authors that might negatively impact their ability or willingness to share their data. Collaboration with the developers of PLINK 2.0 (https://www.cog-genomics.org/plink/2.0/) eased submission further as their open source software now supports direct output to GWAS-SSF format. Submitters using other formats have access to a suite of tools for formatting and validation which can be run locally via the command line or through the SSF-morph web interface (https://ebispot.github.io/gwas-sumstats-tools-ssf-morph/). The latter contains a series of pre-generated configuration files supporting instant conversion of common GWAS analysis software outputs, including REGENIE, BOLT-LMM, SNPtest and SAIGE (17-20). Other formats are supported by custom configuration files generated on demand. Metadata submission is made via an Excel form and validated upon submission, and helpdesk support is available from GWAS Catalog staff throughout the process.

Summary statistics metadata defined in the GWAS-SSF standard are presented alongside the data file in .yaml format designed to be both human and computer-readable. Metadata yaml files are now available for all submissions corresponding to the current and previous (21) standard format. Since the two standards (GWAS-SSF and pre-GWAS-SSF) contain different mandatory columns, a label in the YAML file indicates the version. Updates to metadata files - for example, made during QC/annotation - are tracked in the “metadata_last_updated” field. These updates ensure that users have access to key metadata for all summary statistics in an interoperable format, ensuring downstream usability of the data.

Summary statistics are made available in the submitted format and with a harmonised version, in which genomic position data is reported against GRCh38, alleles orientated to the forward strand, and unharmonisable variants (average ∼3.5%) are removed. To support the GWAS-SSF standard and scale to the rate of summary statistics deposition, improvements to our harmonisation pipeline were needed. Enhancements include increased efficiency and portability, allowing users to integrate the pipeline into their own environments. Average processing time per file has been reduced by 75% through three key improvements: use of an efficient Python library - pysam, integrating Nextflow as a workflow manager to efficiently manage resources, and optimizing palindromic variant orientation by sampling 10% of variants in each rather than the entire file (Supplementary Figure 1). The entire body of summary statistics containing the minimum required data fields (i.e. all GWAS-SSF files and 96% of pre-GWAS-SSF files; the remainder are unformatted legacy files or non-standard submissions such as CNVs or gene-based analyses) has now been made available in harmonised format, with new releases daily. Harmonisation output includes a tabix index file for quick data retrieval and a report file detailing the harmonisation reference and process, enabling users to better interpret the harmonised data (Supplementary Figure 2).

The rapid increase in the number of studies in the Catalog in recent years has been driven in large part by large-scale publications (22,23) analysing hundreds to thousands of molecular quantitative traits (protein, lipid and metabolite measurements) across multiple cohorts. These constituted 82% of all studies added to the Catalog in 2023 (Figure 1), and represent a major and expanding set of open data bridging genetic associations to disease/trait mechanisms via proteins and metabolites, as well as enabling identification of potential drug targets (24). Mapping protein and metabolite traits to chemistry ontologies is necessary for integration of GWAS with protein, chemistry and pathway databases and downstream analyses, but this is challenging to do at scale due to differences in the metabolomics platforms and ambiguity of identifiers. We have therefore collaborated with the Ontologies team at EMBL-EBI to map quantitative traits to the Ontology of Biological Attributes (25), where each term contains mapping to chemical entities in CHEBI, and accurate classification of protein or metabolite class - for example, http://purl.obolibrary.org/obo/OBA_2045052. This allows efficient generation of ontology terms at scale, enabling data to be made available more quickly, providing richer annotation for users, and paving the way for future improvements in GWAS Catalog search and visualisation functionality. We encourage authors of large molecular quantitative trait studies to include database identifiers such as CHEBI (26), Protein Ontology (27) or UniProt (28) IDs in their publications and data submissions, to enable accurate trait mapping and maximise reusability of the data.

**Figure 1.**
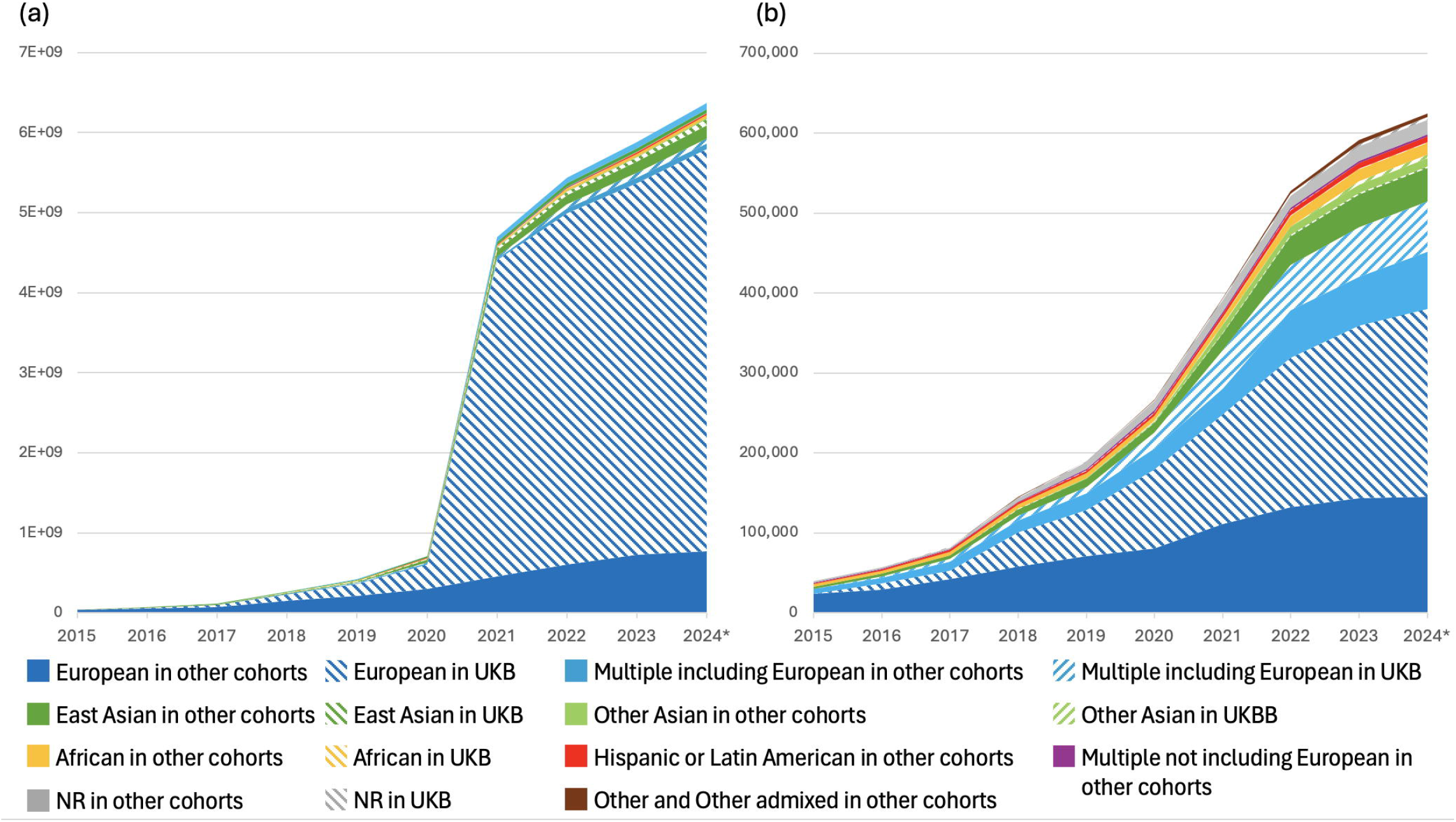
(A) Number of studies added to the GWAS Catalog per year between 2019 and 2024 (until end of August), showing the growing proportion of molecular quantitative trait studies, defined as all studies annotated with the ontology terms “protein measurement” (EFO_0004747), “metabolite measurement” (EFO_0004725), “lipid measurement” (EFO_0004529) or their child terms. (B) Of all studies annotated with these terms, 54% were annotated with a protein term, 32% with a metabolite term and 11% with a lipid term, while 3% included two or more categories.

### UI improvements

The publication rate of GWAS continues to grow and the GWAS Catalog has responded by scaling up our data ingest processes over recent years: in the first half of 2024 we added ∼3,900 studies and ∼8,600 associations per month, compared to 92 studies (42-fold increase) and ∼3,300 associations (2.6-fold increase) per month during the same period of 2020.

The resulting increased data volumes require improved access and visualisation tools, including efficient display of data from large datasets - for example, with thousands of GWAS corresponding to a single publication, thousands of associations for a single variant, or traits with many child terms in the ontology, such as “protein measurement”. Page loading times are improved by up to 95% for these studies due to enhancements in the GUI (Figure 2A). Access to summary statistics is more intuitive and visible, studies with summary statistics in a Publication or Trait page are listed in a separate tab, and clearer indication of whether specific studies have summary statistics or not is provided in the GUI and flat file downloads.

**Figure 2.**
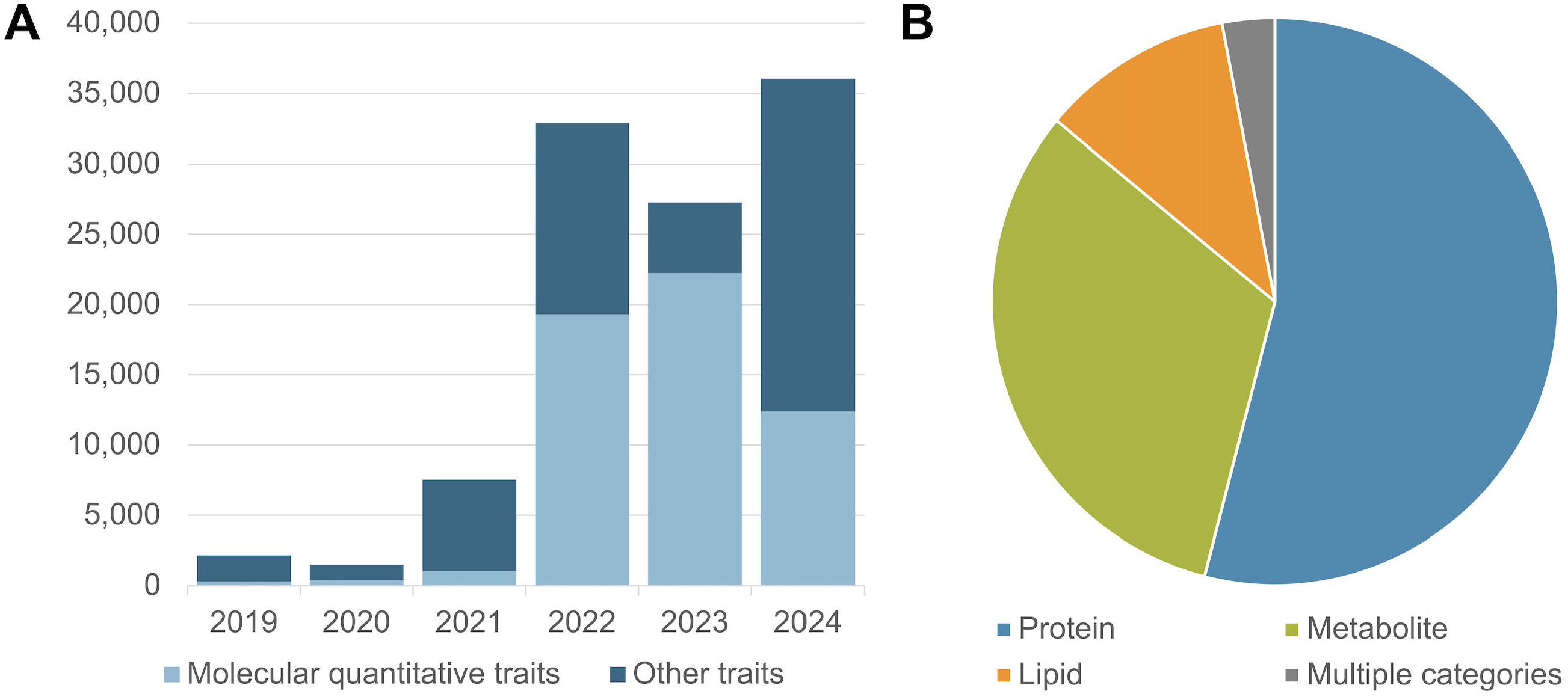
**(A)** New layout of GWAS Catalog UI trait page for “lung cancer”. Data is loaded into each tab (e.g. Full summary statistics) only when users click, to improve performance and load times [1]. Improved pagination allows rows to be loaded in batches, with the user able to choose how many rows to display [2]. If required, pages with many rows can be downloaded for analysis offline without having to load the full dataset in the browser. **(B)** New page displaying all studies in the Catalog, with checkboxes to filter the table to show only specific study types (currently GxE or seqGWAS studies). **(C)** The Study Information panel on each Study page now includes a flag to indicate if it is a GxE study.

We have also improved the visibility of GxE association studies investigating interactions between genetics and environmental exposures such as diet, social inequalities, pollution and infectious agents. GxE data in the GWAS Catalog exceeds 850 studies and 10K associations as of 1st July 2024. The presence of a GxE interaction term in a study is clearly indicated on the study page (Figure 2C), and in the studies download file. In a new page (https://www.ebi.ac.uk/gwas/studies/), users can filter the entire list of studies to show only GxE interactions (Figure 2B). Interest in GxE data is anticipated to grow as the scale of exposome data expands, enabled through increased documentation of external exposure parameters (e.g. air pollution from satellite data), the use of wearables and the adoption of - omics techniques that indicate either proxies (e.g. DNA methylation as an indicator of smoking) or direct measures of exposures (e.g. metabolomics of chemical pollutants) (29). Improved findability in the GWAS Catalog will facilitate the re-use and integration of GxE data as the research area develops further.

Users can also filter for studies exclusively based on whole genome/exome sequencing (seqGWAS; 16K studies to date) (Figure 2B). In future, we will expand this search functionality to include other data types such as CNV- and gene-based association studies, as more become available.

### Population descriptors

Population structure, meaning presence of subgroups within a population, can be attributed to sampling biases, geographic origin, and historical demographic events. If not adequately controlled for, this substructure can confound GWAS producing associations that reflect sample stratification rather than genetic effects on the trait of interest, thus impacting downstream analyses. Population stratification is often controlled for by performing the analysis in a genetically homogenous population where the genetic background of the individuals is described based on similarity to a reference panel or using self-reported ancestry as a proxy for this. This information is also used downstream when choosing an appropriate linkage disequilibrium matrix for fine mapping or in matching a target population to apply a PGS.

The Catalog’s ancestry framework (8) records the author’s sample annotation and also structures population descriptors into broader categories to enhance findability. More than 250 different population terms have been curated in the Catalog, which vary from self-reported ancestry backgrounds to ethnicity, geographical ancestry, founder population or genetic similarity to a reference panel. The addition of the broad ancestry category label enhances the search and usability of the data, making it less time-consuming and computationally less intensive. A clear example of the utility of GWAS Catalog population data can be seen in the GWAS Diversity Monitor, which provides quasi-real-time monitoring of trait and ancestry label diversity to highlight gaps (30).

Whilst the accurate description of populations under study remains important to the reusability of GWAS data, concerns have been raised in recent years (31,32) over the misappropriation of genetic data to support typological thinking around race and ancestry, and the misconception of race as a biological attribute. These were addressed in a 2023 NASEM report (33) which made a number of important recommendations on language relevant to GWAS including: explaining why the use of population descriptors is necessary in the context of the research presented; describing the population in terms of genetic similarity to a reference panel wherever possible, e.g. “1000 Genomes YRI”; avoiding the use of the term “ancestry” where ancestry per se has not been studied; acknowledging that any population grouping is essentially arbitrary, and where group labels are used they should be described as such; and avoiding typological thinking. We have therefore updated the language on our website and in our documentation (https://www.ebi.ac.uk/gwas/docs/ancestry) to clarify the rationale for using population descriptors. Use of “ancestry” remains common in the genomics literature so we have retained the term but provided a clearer explanation and de-emphasised its use when possible - for example, the relevant documentation page is now called “Population descriptors”, https://www.ebi.ac.uk/gwas/populationdescriptors. We have changed the term “ancestry category” to “ancestry label” to make clear that this label is applied to enable search and not as a biological or scientific category. To support the move towards genetic similarity to a reference panel, we have piloted the extraction of reference panels from GWAS publications but found that this is not yet being reported for the majority of GWAS samples (Supplementary Figure 3). We will work towards the inclusion of reference panel information in the GWAS Catalog in the future. In addition, we request summary statistics submitters to provide additional sample metadata such as the method used to define the population descriptor provided (self-reported/genetic similarity), and the imputation panel used. Cohort names are now curated when provided, aligning with the requirement for greater transparency in sample description. Cohorts are now available in the studies download file for all studies curated since the pilot work to extract this information began in 2020. Cohort abbreviations from the discovery stage GWAS are either extracted from the literature or supplied by submitters to match a predefined list shared with the PGS Catalog (https://ftp.ebi.ac.uk/pub/databases/spot/pgs/metadata/pgs_all_metadata_cohorts.csv), seeded from Mills & Rahal, 2019 (30). Consortium names are extracted if component cohort names are absent, or there are too many component cohorts to efficiently curate. Analysis of GWAS-cohort association indicates that UK Biobank (UKB) remains the most common cohort at 13% of publications and represents 30,800 GWAS, likely due to deep phenotyping and ease of data access, followed by FinnGen (∼2% since 2020) and BioBank Japan with (∼1.3%). Provision of cohort annotation allows GWAS Catalog users to account for or avoid sample overlap in downstream analyses. UKB GWAS data is typically restricted to “White” or “White British” samples, and this contributes significantly to the European bias in GWAS results (Figure 3). 71% of publications and 83% of studies where UKB data is part of the analysis include only samples with the European ancestry label. While the panUKB (34) multi-ancestry analysis of ∼7K traits in UKB is in preprint at the time of writing, and future inclusion of these data will contribute to addressing the balance, the general trend of authors performing European-only analyses persists.

**Figure 3.**
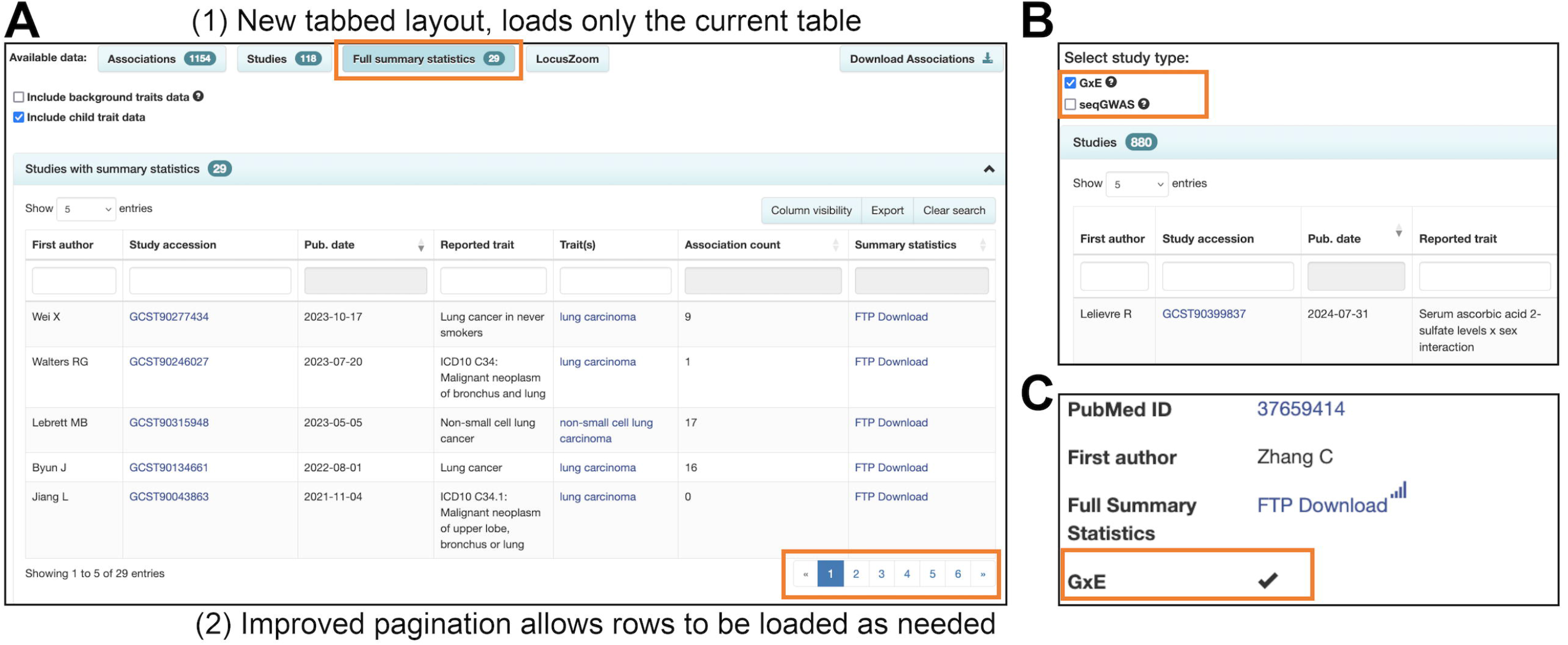
Contribution of individuals from UK Biobank (UKB) to the breakdown by ancestry label of (a) individuals and (b) associations in the GWAS Catalog. The effect on publications and studies can be found in Supplementary Figure 4.

## Discussion

The GWAS Catalog has provided curated GWAS data for 15 years and received deposited summary statistics for the last 5. FAIR human GWAS data from the Catalog is widely used in translational research such as the generation of polygenic scores, Mendelian randomisation, and in drug discovery pipelines. Considering the growth in GWAS data volume and GWAS Catalog content, programmatic access is an increasingly important route. Future software releases will include a new version of the GWAS Catalog REST API (our existing API will be retained), with a more simplified and HATEOAS compliant JSON response structure, full search capability comparable to the web interface, and improved compatibility with downstream software packages such as gwasrapidd (35). Shared submission infrastructure with the PGS Catalog will simplify sharing of GWAS and PGS together and improve data completeness.

Absence of GWAS reflecting global genomic diversity limits the translational potential of GWAS data to benefit all populations. However, generation and access to multi-ancestry data requires funding and infrastructure. Therefore, the GWAS Catalog has been working with researchers from low- and middle-income countries to provide training, support data sharing and remove barriers to data access and use. Over-representation of samples similar to European reference populations also derives from selection of cohorts by researchers, and where multi-ancestry components exist these may be excluded at the point of analysis or publication due to power concerns. For example, out of ∼500K individuals in UKB, 30K from minority groups are frequently excluded from publications (36). It is therefore important to ensure FAIR descriptions of populations allowing monitoring of inclusion at all levels of the study cycle, whilst avoiding language that bolsters racist misappropriation of genetics and genomics research. This is a principal discussion point for the community in the near future. GWAS authors and submitters are encouraged to use genetic similarity measures to define population groups and state any reference panels used in unambiguous language, enabling data resources such as the GWAS Catalog to provide accurate and interoperable data. To increase the reproducibility of GWAS findings and power to discover novel loci, diverse samples are required not only in terms of ethnicity or ancestry but also in environmental and demographic variables, which have been shown to have substantial effects, for example, in the transferability of polygenic scores (37-39). Describing the environment of samples in a consistent way is a challenge for the FAIRification of data. It requires community consensus on reporting - for example, by using geographic location data that can be linked to databases of environmental measurements and indices. Definition of standard cohort descriptors will provide a key component of this effort by enabling text mining and linking between data sources. Consistency is currently lacking - for example, samples from the MyCode Community Health Initiative of the Geisinger Health System are described as MyCode, Geisinger, GHS and other variants. We are working with other databases towards creating a shared resource to facilitate linking datasets from the same cohort that exist in different databases.

The future of GWAS studies will likely be driven by increased availability of large, deeply phenotyped whole genome sequencing datasets, allowing genome-wide differences in structural and copy number variation (CNV-SV) to be systematically evaluated at scale. Such variants are well-known to functionally affect human traits, and represent important potential therapeutic targets. Development of standards in CNV-SV association reporting will be critical to fully integrate data into repositories such as the GWAS Catalog and serve it alongside SNP association data (40) and maximise its utility in functional analyses. By continuous engagement with the human genetics community, we defined a standard format for summary statistics and developed tools to facilitate the submission and reusability of the data in meta-analysis, generation of polygenic scores, Mendelian randomisation and other downstream uses. We continue to work with the community, via workshops, conferences and surveys, on additional data content and structure with the most benefit for human genetics research.

## Data availability

The GWAS Catalog is an open-source project and code is available in the project’s github repository (https://github.com/EBISPOT/goci). Curated data are available from the query interface (https://www.ebi.ac.uk/gwas/) and download files from https://www.ebi.ac.uk/gwas/downloads. APIs for the summary statistics data (http://www.ebi.ac.uk/gwas/summary-statistics/api/) and the curated data (https://www.ebi.ac.uk/gwas/docs/programmatic-access) provide programmatic access to all the Catalog’s data. The Catalog’s GUI also provides access to summary statistics https://www.ebi.ac.uk/gwas/downloads/summary-statistics. GWAS summary statistics submitted after March 2021 are made available under CC0 terms (https://creativecommons.org/publicdomain/zero/1.0/), while those submitted prior to March 2021 are made available under the standard terms of use for EBI services (https://www.ebi.ac.uk/about/terms-of-use/). We advise consumers of data hosted by the GWAS Catalog to note the licence terms of individual datasets. Other GWAS Catalog data is covered by the EBI terms of use. Code is available under the Apache version 2.0 licence (http://www.apache.org/licenses/LICENSE-2.0).

## Funding

National Human Genome Research Institute of the National Institutes of Health: [1U24HG012542-01; ‘Phenomics First’ RM1 HG010860 (to J.M., R.S., A.I.)]

Open Targets*: [OTAR2045, EBI-OT02, EBI-OT01]

Office of the Director, National Institutes of Health [R24 OD011883] (to J.M., R.S., A.I.) Human Ecosystems Transversal Theme (European Molecular Biology Laboratory) European Molecular Biology Laboratory Core Funds

National Institute of Diabetes and Digestive and Kidney Diseases [UM1DK105554] Munz Chair of Cardiovascular Prediction and Prevention (to M.I.)

UK Economic and Social Research 878 Council [ES/T013192/1] (to M.I.).

British Heart Foundation core funding [RG/18/13/33946: RG/F/23/110103] (to M.I., S.A.L.)

NIHR Cambridge Biomedical Research Centre [NIHR203312] (to M.I., S.A.L.), BHF Chair Award [CH/12/2/29428] (to M.I., S.A.L.)

Health Data Research UK** (to M.I., S.A.L.)

Funding for open access charge: NIH grant 1U24HG012542-01 and/or EMBL core funds. J.M.. and P.H. are employees of the National Human Genome Research Institute.

*Open Targets is a pre-competitive collaboration between EMBL-EBI, Genentech, GSK, Pfizer, Sanofi, Takeda, and the Wellcome Trust Sanger Institute.

**Health Data Research UK is funded by the UK Medical Research Council, Engineering and Physical Sciences Research Council, Economic and Social Research Council, Department of Health and Social Care (England), Chief Scientist Office of the Scottish Government Health and Social Care Directorates, Health and Social Care Research and Development Division (Welsh Government), Public Health Agency (Northern Ireland), British Heart Foundation and the Wellcome Trust.

Conflict of Interest statement: M.I. is a trustee of the Public Health Genomics (PHG) Foundation, a member of the Scientific Advisory Board of Open Targets, and has research collaborations with AstraZeneca, Nightingale Health and Pfizer which are unrelated to this study. No other conflicts of interest were reported by authors.

## Acknowledgements

The authors thank the GWAS Catalog’s users, study authors, submitters of summary statistics and our Scientific Advisory Board. We also thank the EMBL-EBI Technical Services Cluster for maintenance of the computational infrastructure, the EMBL-EBI Training team for supporting provision of our training materials, and the many GWAS experts who have contributed to discussions about Catalog functionality, data representation and user experience testing. We acknowledge the insight of Teri Manolio, Lucia Hindorff and Leah Mechanic, which has informed the Catalog’s development. The content is solely the responsibility of the authors and does not necessarily represent the official views of the National Institutes of Health, the NIHR or the Department of Health and Social Care.

