## Supplementary Information for "The NHGRI-EBI GWAS Catalog: standards for reusability, sustainability and diversity"

### Supplementary Figure 1

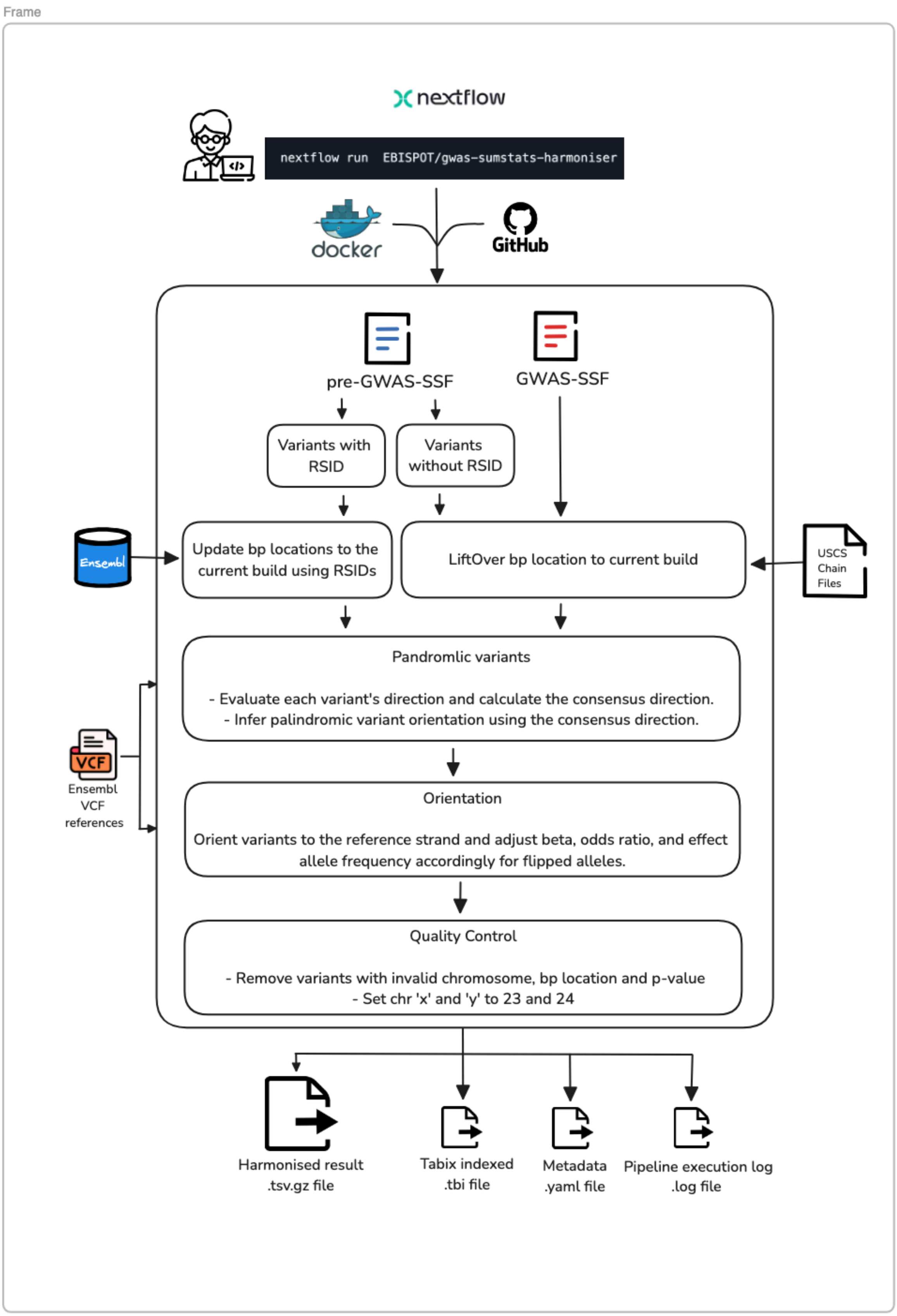


**SFigure 1**. Flow diagram illustrating the GWAS Catalog's summary statistics harmonisation process.

### Supplementary Figure 2

################################################################

**HARMONISATION RUNNING REPORT**

################################################################

**1. Pipeline details**

**A. Pipeline Version: 0.1.0**

**B. Running date: Jul 29 2024**

**C. Input file: GCST90428625.tsv.gz**

################################################################

**2. Reference data**

**##source=ensembl;version=95;url=http://vertebrates.ensembl.org/homo_sapiens**

**##reference=ftp://ftp.ensembl.org/pub/release-95/fasta/homo_sapiens/dna/**

**##ID=dbSNP_151,Number=0,Type=Flag,Description="Variants (including SNPs and indels) imported from dbSNP"**

################################################################

**3. Mapping result**

**0.0465071% (3777 sites out of 8121338) were dropped because they could not be mapped.**
**99.9535% (8117561 sites) were carried forward.**

################################################################

**4. Palindromic SNPs**

**palin_mode: forward**

**Direction of palindromic SNPs inferred as forward by establishing consensus direction of 10% of all sites (forward sites ratio =0.9991262896676848).**

################################################################

**5. Successfully harmonised variants**

**99.77% ( 8098600 of 8117561 ) sites successfully harmonised.**

| hm_code | Number | Percentage | Explanation |
|---|---|---|---|
| 10 | 6858838 | 84.49% | Forward strand; Correct orientation; Already harmonised |
| 11 | 10277 | 0.13% | Forward strand; Flipped orientation; Requires harmonisation |
| 12 | 5948 | 0.07% | Reverse strand; Correct orientation; Already harmonised |
| 13 | 226 | 0.00% | Reverse strand; Flipped orientation; Requires harmonisation |
| 5 | 1220306 | 15.03% | Palindromic; Assume forward strand; Correct orientation; Already harmonised |
| 6 | 3005 | 0.04% | Palindromic; Assume forward strand; Flipped orientation; Requires harmonisation |

################################################################

**6. Failed harmonisation**

**0.23% ( 18961 of 8117561 ) sites failed to harmonise.**

| hm_code | Number | Percentage | Explanation |
|---|---|---|---|
| 15 | 18961 | 0.23% | No matching variants in reference VCF; Cannot harmonise |

################################################################

**7. Overview**

| | |
|---|---|
| Result | SUCCESS_HARMONIZATION |

**SFigure 2.** Example of a harmonisation report file. The file summarises the reference VCF file used in harmonisation, details of the process used to orientate palindromic variants and percentage of variants harmonised, failed or dropped.

**Supplementary Figure 3**

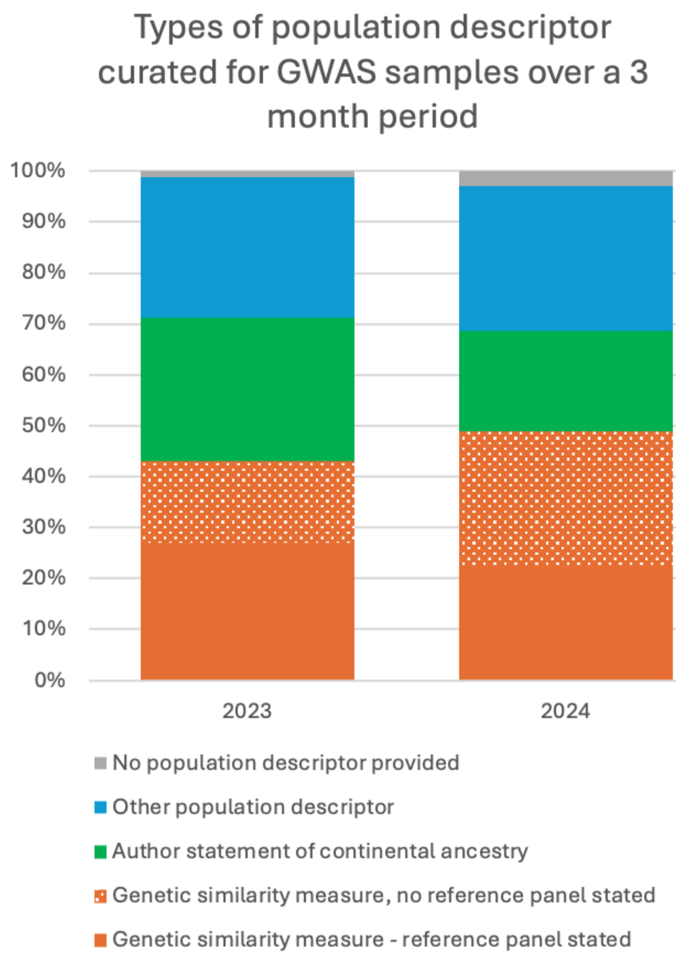


**SFigure 3**. Population descriptors curated for 355 GWAS samples. Over a 10-week period between July and September, GWAS Catalog curators recorded the different types of population descriptor reported by authors which were later used to assign an ancestry category label. This comprised 161 distinct sample groups from 89 GWAS publications in 2023, and 194 distinct sample groups from 119 GWAS publications in 2024. For more details on the methodology, please refer to Morales et al 2018.

Population descriptor types were assigned as follows:

1. Genetic similarity measure, no reference panel e.g. “All participants were of East Asian ancestry based on application of on genetic principal component analysis (PCA) to identify population stratification outliers”

2. Genetic similarity measure, reference panel stated e.g. “We used 1000G Yoruba (YRI, n=108) as the reference population for the African ancestry”

3. Author statement of continental ancestry with no information as to ascertainment method e.g. “All participants were of European ancestry”

4. Other population descriptors, such as self-reported continental ancestry (e.g. all participants self-reported as European ancestry”), ethnicity (e.g. “Participants were Han Chinese”), country of recruitment (e.g. “Participants were recruited in Iceland”) and others (e.g. the Harmonised Race and Ethnicity groups assigned by Million Veteran Program).
5. No curatable population descriptor provided.

**Supplementary Figure 4**

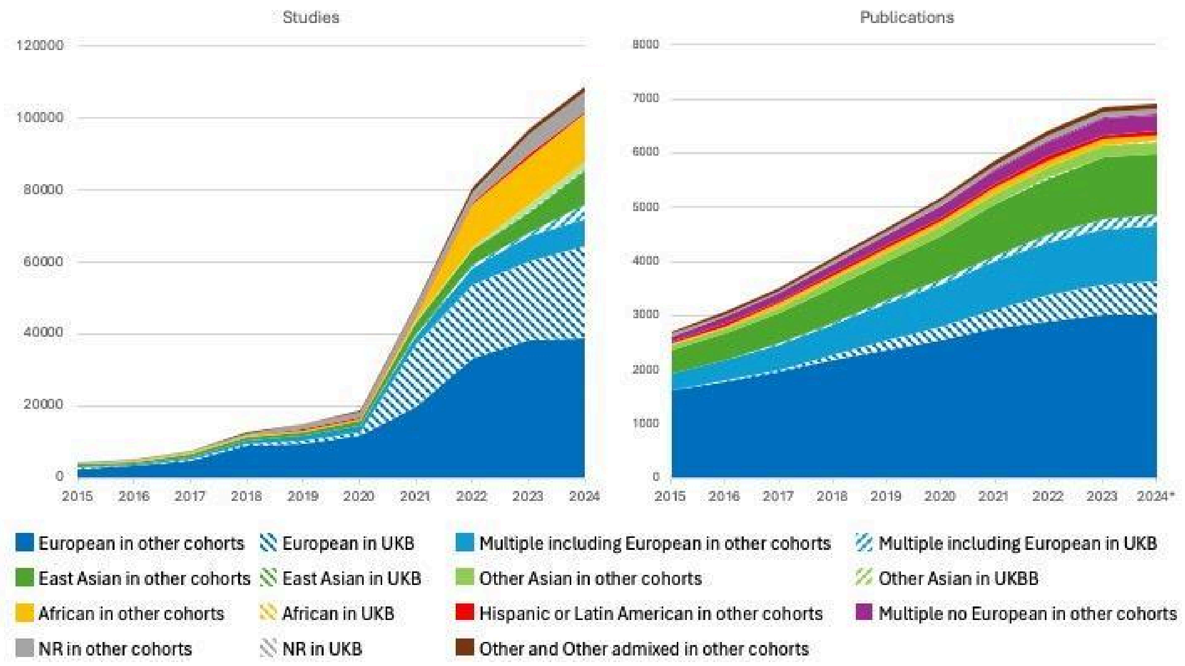


**SFigure 4**. Contribution of individuals from UK Biobank (UKB) to the breakdown by ancestry label of (a) studies and (b) publications in the GWAS Catalog.
