## Supplementary figures and images for "The NHGRI-EBI GWAS Catalog: standards for reusability, sustainability and diversity"

### Supplementary Figure 4

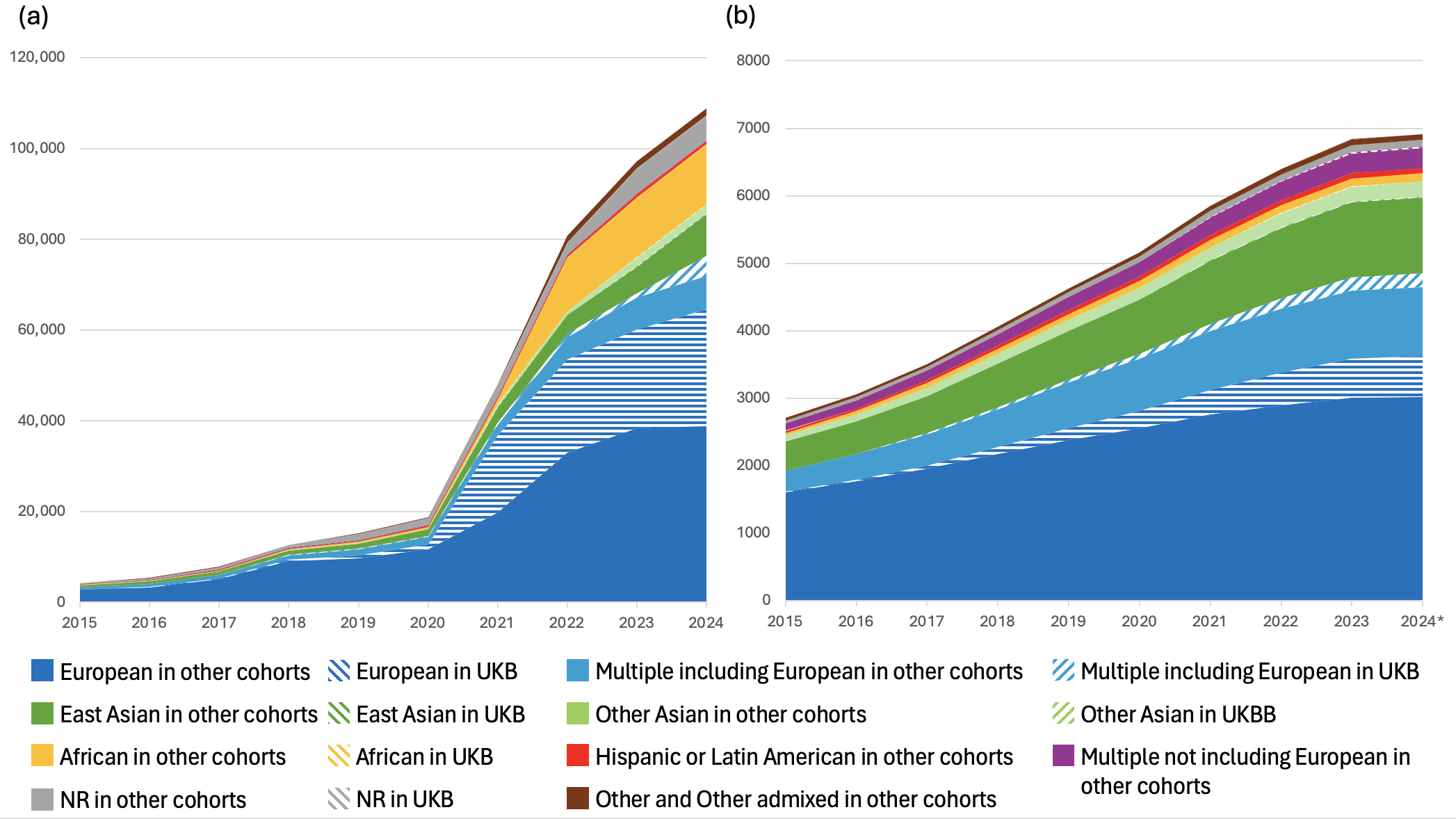
